# Astrocyte- and neuron-derived extracellular vesicles from Alzheimer’s disease patients effect complement-mediated neurotoxicity

**DOI:** 10.1101/2020.04.14.041863

**Authors:** Carlos J Nogueras-Ortiz, Vasiliki Mahairaki, Francheska Delgado-Peraza, Debamitra Das, Konstantinos Avgerinos, Matthew Hentschel, Edward J Goetzl, Mark P Mattson, Dimitrios Kapogiannis

## Abstract

We have previously shown that blood astrocytic-origin extracellular vesicles (AEVs) from Alzheimer’s disease (AD) patients contain high complement levels. To test the hypothesis that circulating EVs from AD patients can induce complement-mediated neurodegeneration, we assessed the neurotoxicity of immunocaptured AEVs (with anti-GLAST antibody), neuronal-origin NEVs (with anti-L1CAM antibody), and multicellular-origin (with anti-CD81 antibody) EVs from the plasma of AD, frontotemporal lobar degeneration (FTLD) and control participants. AEVs (and, less effectively, NEVs) of AD participants induced Membrane Attack Complex (MAC) expression on recipient neurons, membrane disruption, reduced neurite density, and decreased cell viability in rat cortical neurons and human IPSC-derived neurons. Neurodegenerative effects were not produced by multicellular-origin EVs from AD participants or AEVs/NEVs from FTLD or control participants, and were suppressed by the MAC inhibitor CD59 and other complement inhibitors. Our results support the stated hypothesis and suggest that neuronal MAC deposition is necessary for AEV/NEV-mediated neurodegeneration in AD.

## Introduction

Alzheimer’s disease (AD) is the result of a neurodegenerative cascade involving progressive deposition of misfolded amyloid beta-peptide (Aβ) and tau, abnormalities in brain cell homeostasis and function, and ineffective or even maladaptive compensatory mechanisms. Recently, neuroinflammation and its cellular mediators, microglia and astrocytes, have emerged as important factors in AD pathogenesis (1) and have been implicated in the development of both Aβ (2, 3) and tau (4) pathologies. Maladaptive neuroinflammation in AD involves the complement cascade (5, 6), a system of sequentially activated humoral and cellular proteins that lies on the interface between innate and adaptive immunity and promotes host defenses against infection and tissue homeostasis. The complement cascade can be activated by IgM or IgG immunocomplexes (classical pathway), bacteria or toxins (alternative pathway) or mannose residues (lectin pathway) (7), ultimately leading to the formation of the membrane attack complex (MAC), a cytolytic membrane pore formed by the sequential assembly of soluble complement proteins C5b, C6, C7, C8 and C9, that causes cell osmolysis after intercalation into the plasma membrane (8, 9). For peripheral tissues, the main source of complement proteins is the liver, whereas the brain relies on local synthesis observed in all cell types (10–12). Appropriately regulated complement is necessary for brain defense and homeostasis, but overactivated complement may lead to brain pathology.

Multiple lines of evidence implicate the complement cascade in AD pathogenesis. In AD mouse models, molecular inhibition or genetic deletion of complement factors/receptors, such as C3/C3R, C5/C5R and C1q decrease glial activation, Aβ and hyperphosphorylated tau deposition and synaptic pruning (6, 13–15). Complement components of the classical pathway colocalize with Aβ plaques and tau neurofibrillary tangles in the hippocampus, temporal and frontal lobes of AD patients (12, 16–18). In addition, the MAC has been detected in the vicinity of Aβ and tau deposits, and both in the inner and outer leaflets of the neuronal plasma membrane (19). Furthermore, genome wide association studies (GWAS) have associated CR1 (gene encoding the receptor for C3b/4b) with late onset AD (20); the interaction between C3b and CR1 mediates Aβ phagocytosis (21); thus polymorphisms altering CR1 function could decrease Aβ clearance.

Extracellular vesicles (EVs) are membranous nanoparticles produced by all cells including neurons and astrocytes. The role of EVs in the brain is multifaceted and includes the exchange of cargo molecules between neurons and glia to regulate neuronal activity (22), but also the transfer of misfolded α-synuclein, Aβ, and hyperphosphorylated tau, possibly contributing to neurodegenerative disease propagation (23, 24). We and others have isolated neuronal and astrocytic origin-enriched EVs (NEVs and AEVs, respectively) from plasma and human cells by particle precipitation followed by immunocapture with anti-GLAST and anti-L1CAM antibodies respectively and have shown that they are carriers of AD pathogenic proteins (25–27). Recently, we showed that AEVs of AD patients carry high levels of multiple complement components of the classical and alternative pathways (i.e. C1q, C4b, C3b, Bb, C3d, factor B, factor D), as well as the MAC (28). Interestingly, we also found levels of endogenous complement regulatory proteins, such as CD59, that binds to the transmembrane residues C8 and C9 and blocks the formation of the MAC (8, 9), CD46 and the decay-accelerating factor (DAF), to be low in EVs from AD patients (28).

Based on these findings, we hypothesized that neuronal uptake of AEVs may lead to delivery and/or *in situ* formation of MAC, and ensuing neurodegeneration. In the present study, we confirm this hypothesis by showing that circulating AEVs and NEVs of AD patients are indeed neurotoxic through MAC deposition, membrane disruption, and necroptosis, findings that open new avenues for therapeutic intervention in AD.

## Results

### Characterization of plasma-derived EVs

According to established criteria (29), we characterized our EV preparations by determining the size and concentration of isolated EVs by Nanoparticle Tracking Analysis (NTA) (Fig. S1a) and Transmission Electron Microscopy (Fig. S1c): the size distribution of nanoparticles immunocaptured by L1CAM, GLAST or CD81 was consistent with a mixed EV population likely predominated by exosomes (50-100 nm) and microvesicles (100-200 nm), whereas total EVs showed higher diameters consistent with co-sedimentation of EVs and vesicle-like lipoproteins (Fig. S1b). EV purification of immunoprecipitated EVs was confirmed by western blots showing enrichment for transmembrane and intravesicular EV markers (i.e. CD81 and alix), low levels of apolipoprotein A1 and absence of the Golgi-specific negative EV marker GM-130, in comparison with EV-depleted plasma and total EVs, as previously seen (27) (Fig. S1c).

### AD AEVs and NEVs are neurotoxic

EVs from individual subjects were not pooled and their effects were assessed and analyzed separately to respect their biological variability. First, we sought to determine whether AEVs labelled with the fluorescent lipid analogue PKH26 may be internalized by cultured E18 rat cortical neurons (SI Appendix, Methods; Fig. S2), as previously shown for NEVs (30). After 1 hour of incubation, we detected puncta-like PKH26+ structures in the soma and neurites of cells treated with AEVs of AD and normal participants (Fig. S2c). Then, we used the MTT cell viability assay to evaluate whether AEVs and NEVs from participants with AD (n = 7) compared to cognitively normal controls (n = 6) may be neurotoxic in a concentration- and time-dependent manner (Fig. 1). Both AEVs and NEVs from AD compared to control participants decreased cell viability, with maximum effects produced by a concentration of 600 EVs/neuron (Fig. 1a) and incubation for 48 hours (Fig. 1b), treatment conditions that were kept constant in subsequent studies unless otherwise indicated. The neurotoxicity of AD AEVs and NEVs was comparable to excitotoxicity by 10 μM glutamate, hence physiologically relevant (31), and was cell of origin-specific (astrocytic and neuronal) and independent of putative soluble co-precipitates, since no cell viability differences were observed in neurons treated with multicellular-origin CD81+ and total EVs from the same AD participants (Fig. 1c). For NEVs, Group, F (1, 34.703) = 4.544, p = 0.04; AD vs. Normal, p = 0.04. For AEVs, Group, F (1, 39.737) = 10.86, p = 0.002; AD vs. Normal, p = 0.002. For CD81 + EVs, Group, F (1, 26.817) = 0, p = 0.991; AD vs. Normal, p = 0.991. For total EVs, Group, F (1, 30.154) = 0.291, p = 0.593; AD vs. Normal, p = 0.593. In a Model including all EV types and the Group*EV type interaction term, NEVs produced a significant MTT decrease (F (1, 144.874) = 9.151, p = 0.003; AD vs. Normal, p = 0.003), and so did AEVs (F (1, 146.807) = 7.7, p = 0.006; AD vs. Normal, p = 0.006), but CD81 +EVs did not (F (1, 147.225) = 0.015, p = 0.904; AD vs. Normal, p = 0.904) and neither did total EVs (F (1, 147.077) = 0.177, p = 0.675; AD vs. Normal, p = 0.675). For normal controls, there were no significant differences between EV types and no effects on MTT were seen. For AD participants, NEVs and AEVs had similar effects on MTT (p = 0.695), but AEVs decreased MTT compared to both CD81 + (p = 0.035) and total EVs (p = 0.029), and NEVs showed similar strong trends in pairwise comparisons (p = 0.087 and p = 0.067 respectively). Results were similar for models including age and sex. Moreover, there were no effects on the viability of neurons treated with AEVs from 2 participants with frontotemporal lobar degeneration (FTLD) compared to 6 controls (Group, F (1, 35.425) = 0.007, p = 0.934), suggesting that AEV neurotoxicity is not a common feature of neurodegenerative proteinopathies (Fig. 1d).

**Figure 1.**
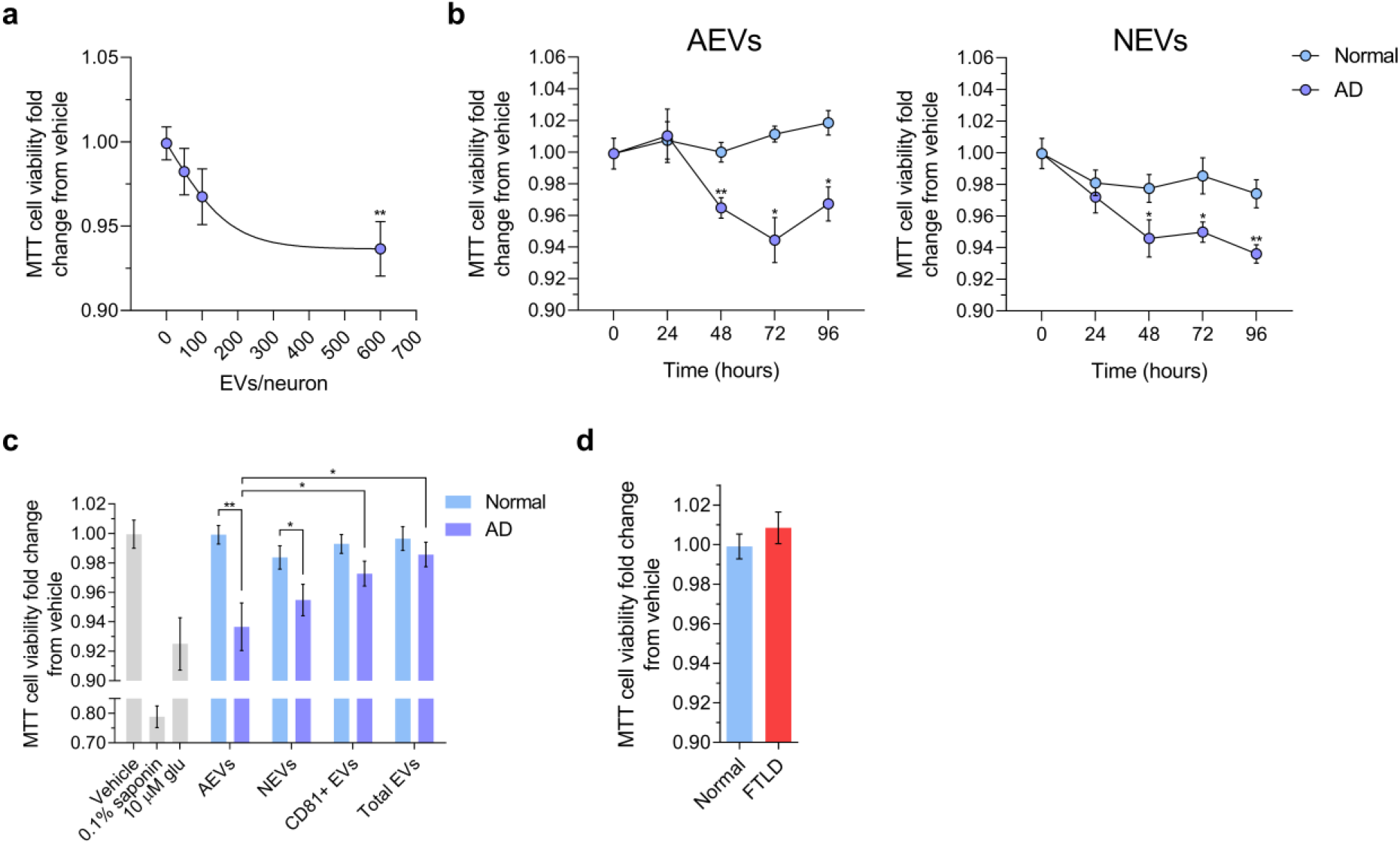
AD AEVs and NEVs impair neuronal viability. **a,** Scatter dot plot showing the MTT cell viability fold change from vehicle of E18 rat cortical neurons treated with AD AEVs in a concentration-dependent manner. Each dot represents the mean value ± the standard error obtained from neurons treated with plasma derived AEVs from 7 AD participants at the indicated concentration or vehicle (0 EVs/neuron) in triplicate. Trendline shows the four-parameter logistic nonlinear regression. **p = 0.0015, 600 vs 0 EVs/neuron; one-way ANOVA corrected for multiple comparisons using the Dunnet test. **b,** Scatter dot plots with connecting line showing the MTT cell viability fold change from vehicle of E18 rat cortical neurons treated in triplicate with AEVs or NEVs from 7 AD participants and 6 normal controls at 600 EVs/neuron in a time-dependent manner. Each dot represents the mean value ± SEM. AD vs Normal: AEVs, 48 h **p = 0.0015, 72 h *p = 0.0191, 96 h *p = 0.0201; NEVs, 48 h *p = 0.0337, 72 h *p = 0.0127, 96 h **p = 0.0016; two-tailed unpaired t-test with a confidence interval of 95%. Time-dependent EV treatment vs vehicle treatment (0 hours): AEVs, 48 h **p = 0.0060, 72 h **** p < 0.0001, 96 h **p = 0.0023; NEVs, 48 h ****p < 0.0001, 72 h *p = 0.0186; 96 h **p = 0.0028; ordinary one-way ANOVA. **c,** Bar graph showing a significant decrease in MTT cell viability (fold change from vehicle) in E18 rat cortical neurons treated with AEVs and NEVs, but not CD81+ EVs and total plasma EVs from 7 AD participants compared to neurons treated with AEVs and NEVs from 6 normal controls respectively. Neurons treated with 0.1% saponin detergent and 10 μM glutamate were used as positive controls for neurotoxicity. Each bar represents the mean value ± SEM from triplicate wells and five different experiments; the condition of 600 EVs/neuron and incubation for 48 hours was selected based on a, b. Analysis was based on mixed linear model accounting for technical (well, experiment) and biological (Subject, Group) variability. For NEVs, factor Group, F (1, 34.703) = 4.544, p = 0.04; AD vs. Normal, p = 0.04. For AEVs, factor Group, F (1, 39.737) = 10.86, p = 0.002; AD vs. Normal, p = 0.002. For CD81 + EVs, factor Group, F (1, 26.817) = 0, p = 0.991; AD vs. Normal, p = 0.991. For total EVs, factor Group, F (1, 30.154) = 0.291, p = 0.593; AD vs. Normal, p = 0.593. *p < 0.05, ** p < 0.01. **d,** Bar graph showing the MTT cell viability fold change from vehicle of neurons treated with AEVs from two FTLD and 6 normal control participants. No difference in cell viability was observed; factor Group, F (1, 35.425) = 0.007, p = 0.934. Treatments were carried out as indicated in **c**.

Furthermore, to assess potential neurodegeneration, we used β III-tubulin fluorescence immunocytochemistry and showed a significant neurite density decrease in neurons treated with AEVs from AD compared to control participants (F (1, 27.290) = 6.113, p = 0.02; AD vs. Normal, p = 0.02); the difference was not observed in NEV-treated (F (1, 23.522) = 0.019, p = 0.891) or CD81+ EV-treated neurons (F (1, 12.803) = 1.440, p = 0.252) (Fig. 2), suggesting a greater neurodegenerative potential for AEVs compared to NEVs or CD81+ EVs.

**Figure 2.**
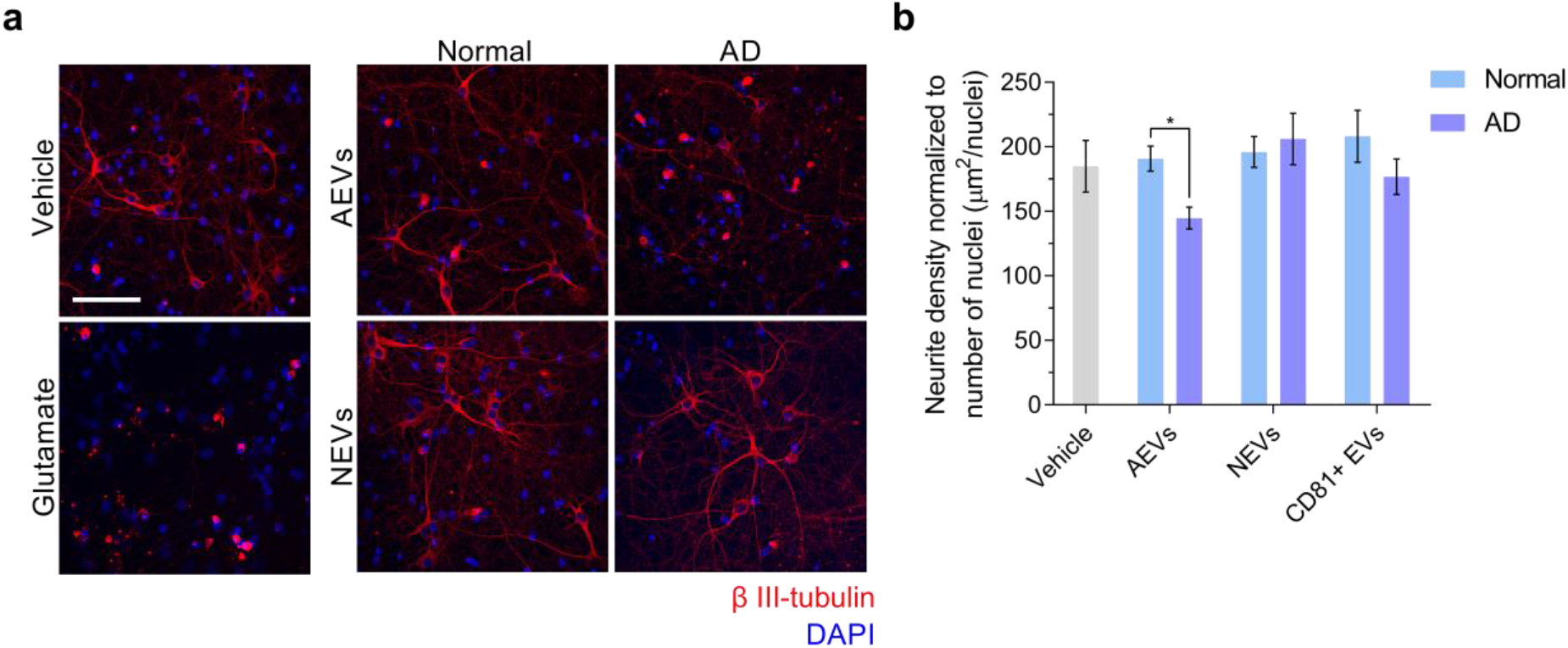
AD AEVs induce neurodegeneration. **a,** Immunofluorescence of β III-tubulin (red) as an indicator of neurite density in E18 rat cortical neurons incubated with either AEVs or NEVs isolated from the plasma of AD patients and normal controls at 600 EVs/neurons for 48 hours. Fluorescent confocal microscopy images at 25X magnification show that neuronal cultures treated with AEVs from AD participants exhibit decreased neurite density compared to vehicle-treated neurons and neurons treated with AEVs from normal participants, as well as with NEVs from AD and normal participants. Treatment with 100 μM glutamate was used as a positive control for neurotoxicity. DAPI staining in blue shows the localization of nuclei. Scale bar, 100 μm. **b,** A bar graph showing the quantification of neurite density defined as the image area covered by β III-tubulin signal above threshold normalized to the number of nuclei. Each bar represents the mean value ± SEM of 5-15 images per treatment with NEVs, AEVs, and CD81+ EVs from 4 AD subjects and 3 normal participants in duplicate. There is significant neurite density decrease in neurons treated with AEVs from AD compared to normal participants (F (1, 27.290) = 6.113, p = 0.02; AD vs. Normal, p = 0.02); the difference was not observed in NEV-treated (F (1, 23.522) = 0.019, p = 0.891) or CD81 + EV-treated neurons (F (1, 12.803) = 1.440, p = 0.252) (Fig. 2), suggesting a greater neurodegenerative potential for AEVs compared to NEVs or CD81+ EVs. *p < 0.05.

### Neurotoxicity of AD AEVs is associated with complement MAC accumulation, membrane disruption and necroptosis activation

Given our previous finding of high levels of complement effectors, including MAC, in circulating AEVs of AD participants (28), we sought to evaluate if treatment with AD AEVs results in MAC deposition and membrane disruption in recipient neurons. First, we performed MAC fluorescence immunocytochemistry in AEV-treated neurons using a validated C5b-9 antibody targeting the C5b-C9 heterodimer, the rate-limiting step of MAC pore formation (9), with β III-tubulin co-staining to visualize neuronal structures. Results showed a dramatic increase of MAC accumulation co-localizing with β III-tubulin in the soma and neurites of cells treated with AD AEVs for 1 hour compared to neurons treated with AEVs from control participants and vehicle-treated [F (1, 2) = 45.527, p = 0.021; AD vs. Normal, p = 0.026; AD vs. vehicle, p = 0.012; Normal vs. vehicle, p = 0.051)] (Fig. 3a, b). To assess whether MAC accumulation leads to plasma membrane disruption, the first step towards osmolysis, we subjected EV-treated cells to the fluorescent ethidium homodimer-1 (EthD-1) membrane disruption assay, which is based on the detection of the membrane impermeable probe EthD-1, which fluoresces in association with nucleic acids (Fig. 3c). Treatment of neurons with NEVs and AEVs from 4 AD compared to 4 control participants resulted in a significantly increased EthD-1 fluorescence fold change over vehicle [For NEVs, F (1, 24.020) = 4.525, p = 0.044; AD vs. Normal, p = 0.044; for AEVs, F (1, 24.872) = 5.804, p = 0.024; AD vs. controls, p = 0.024], suggesting that AD NEVs and AEVs mediate neuronal membrane disruption; there were no differences in EthD-1 fluorescence over vehicle in neurons treated with CD81+ EVs (F (1, 31.429) = 0.969, p = 0.332) from AD compared to control participants. MAC-dependent cytotoxicity is accompanied by activation of the regulated necrosis pathway known as necroptosis, which involves phosphorylation of the mixed-lineage kinase domain-like protein (MLKL) (32, 33). To further explore the mechanism of neurotoxicity induced by AEVs and NEVs, we evaluated whether EV-treated neurons exhibit necroptosis and apoptosis cell death markers. We did not observe activation of apoptosis, as suggested by a fluorescent polycaspase *in vivo* assay that did not show activation of caspases in EV-treated neurons (Fig. 4a, b). This was confirmed by western blots showing that treatment of neurons with AD AEVs does not result in increased levels of the apoptosis markers cleaved caspase-3 and −7 compared to treatment with AEVs from control participants (Fig. 4c, d). Interestingly, neurons treated with AD AEVs showed increased levels of the necroptosis marker phosphorylated MLKL in comparison to control AEVs (Fig. 4c, d).

**Figure 3.**
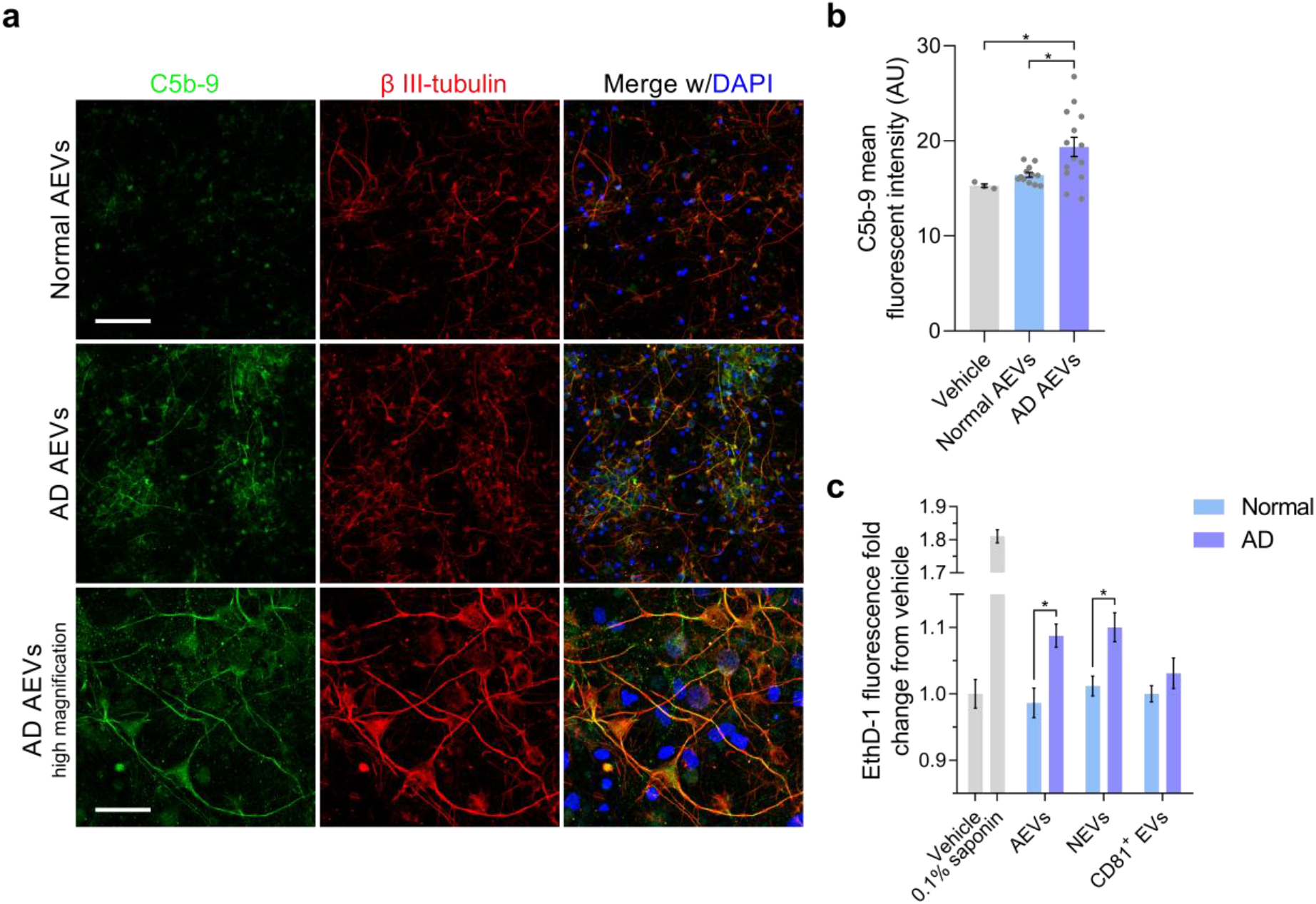
The AD AEV-mediated accumulation of the complement membrane attack complex in neurons is associated with membrane disruption. **a,** E18 rat cortical neurons were incubated with AEVs isolated from the plasma of 2 AD patients and 2 normal controls at 600 EVs/neuron for 1 hour. After incubation, neurons were subjected to C5b-9 immunofluorescence (green), with β III-tubulin (red) co-staining as a marker of neuronal soma and processes. DAPI staining in blue shows the localization of nuclei. Fluorescent confocal microscopy images at 20X magnification show an increased accumulation of the complement membrane attack complex in the neural soma and neurites of cultures treated with AD AEVs in comparison with cells treated with AEVs from normal controls and vehicle [F (1, 2) = 45.527, p = 0.021; AD vs. Normal, *p = 0.026; AD vs. vehicle, *p = 0.012; Normal vs. vehicle, p = 0.051)]. In the bottom panel, an image at 63X clearly shows the neuronal distribution of C5b-9 co-localizing with β III-tubulin. Scale bars: 20X, 100 μm; 63X, 50 μm. **b,** Quantification of **a**. A bar graph with individual values in grey shows the C5b-9 mean fluorescence signal ± SEM obtained from 3-5 images of neuronal cultures treated with AEVs from the plasma of 2 AD patients and 2 normal controls in duplicate. Treatment with vehicle was used to determine the basal detection of C5b-9. **c,** E18 rat cortical neurons were incubated with AEVs, NEVs or CD81 + EVs isolated from the plasma of 4 AD and 4 normal control participants at 600 EVs/neuron for 48 hours. After incubation, neuronal cultures were subjected to the fluorescent EthD-1 membrane disruption assay. Treatment of neurons with NEVs and AEVs from 4 AD compared to 4 normal control participants resulted in a significantly increased EthD-1 fluorescence fold change over vehicle (For NEVs, F (1, 24.020) = 4.525, p = 0.044; AD vs. Normal, p = 0.044); For AEVs, F (1, 24.872) = 5.804, p = 0.024; AD vs. Normal, p = 0.024), suggesting that AD NEVs and AEVs mediate neuronal membrane disruption; there were no differences in EthD-1 fluorescence in neurons treated with CD81 + EVs (F (1, 31.429) = 0.969, p = 0.332) from AD compared to normal control participants. Neurons treated with 0.1% saponin detergent were used as a positive control of neurotoxicity with membrane disruption. Each bar represents the mean value ± SEM of EthD-1 fluorescence fold change from vehicle treatments with EVs from 4 AD and 4 normal control participants in triplicate carried out in three different experiments. *p < 0.05, **p < 0.01.

**Figure 4.**
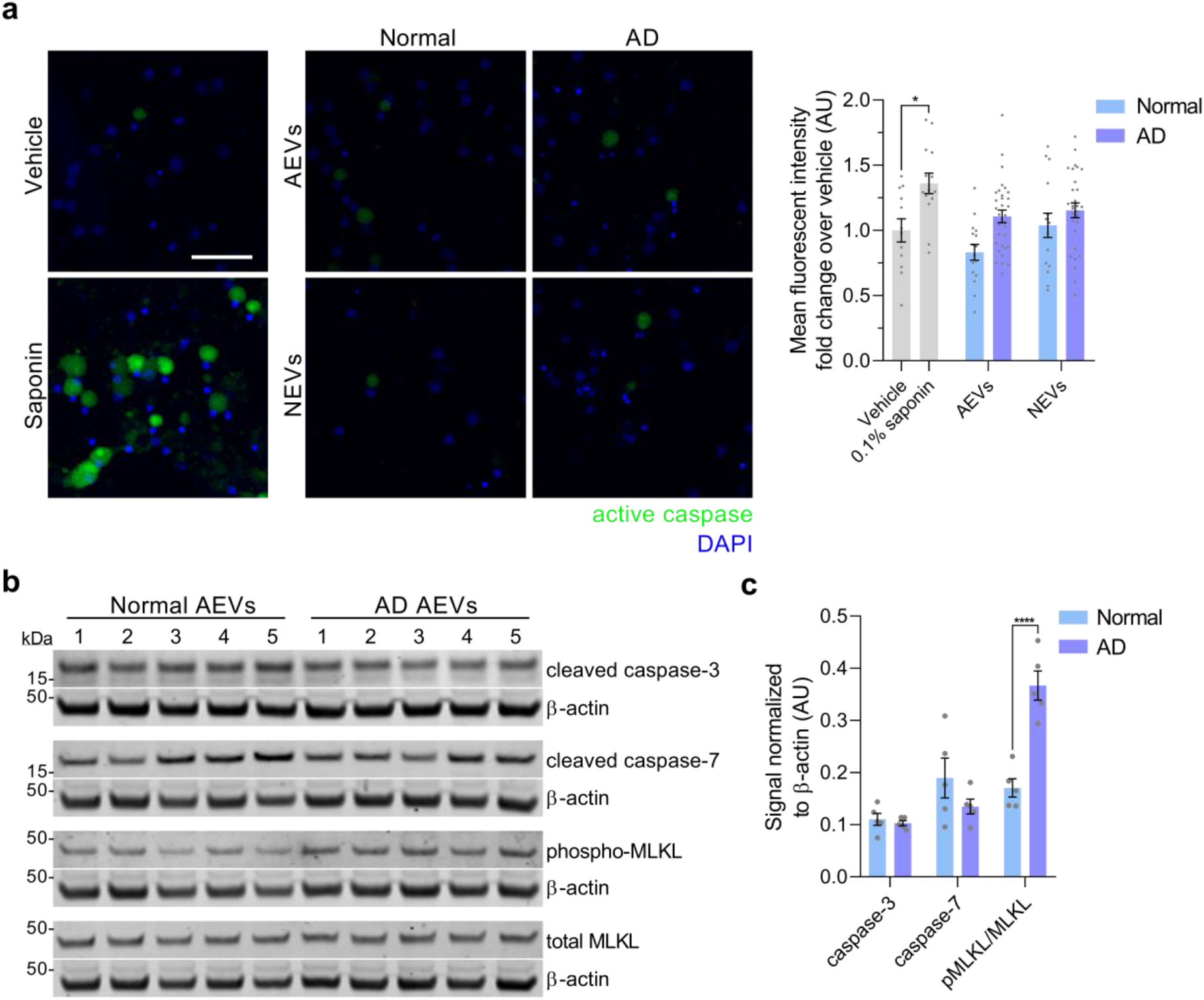
Neuronal activation of necroptosis, but not apoptosis, by AD AEVs. **a,** Neurons treated with AD AEVs were subjected to an *in vivo* fluorescent caspase activity assay that allows the fluorescent detection of active caspases in live cells (green). Treatment with 0.1% saponin detergent was used as a positive control for caspase activation. The membrane permeable nucleic acid dye Hoechst 33342 was used to visualize nuclei (blue). Scale bar, 100 μm. **b,** Quantification of **a.** A bar graph of the quantification of the mean fluorescent intensity in ROIs selected upon thresholding using the Fiji-Image J max entropy algorithm on confocal microscopy images shows that apoptosis is not active in neurons incubated with AEVs and NEVs from 4 AD compared to 4 normal control participants in duplicate. Each bar represents the mean value ± SEM of 5-10 images per treatment. Statistical analysis: one-way ANOVA corrected for multiple comparisons using the Dunnett test; *p = 0.0282. **c,** Western blots against cleaved caspase-3 and −7 as markers of apoptosis, and relative abundance of phosphorylated over total MLKL as a marker of necroptosis. β-actin was used as loading control. **d,** Quantification of **c** showing a significant increase in the relative abundance of phosphorylated MLKL over total MLKL, normalized to β-actin, in rat cortical neurons treated with AD and normal control AEVs. No differences in cleaved caspase-3 and −7 were observed. Each bar represents the mean value ± SEM of the protein optical density normalized to β-actin in neuronal lysates after treatment with plasma-derived EVs from 5 AD and 5 normal control participants in triplicate. Statistical analysis: one-way ANOVA corrected for multiple comparisons using the Dunnett test; ****p < 0.0001.

Next, we replicated our previous finding of low levels of CD59 (a GPI-anchored cell membrane glycoprotein that inhibits MAC assembly and thus protects cells from lysis (8, 9)), in circulating AEVs of AD participants (28) (SI Appendix, Methods section; Fig. S3). Altogether, these findings promote the hypothesis that the neurotoxicity of AD AEVs and NEVs is due to MAC-mediated osmolysis, in an environment impoverished of endogenous complement inhibitors, especially CD59.

### Complement inhibition blocks the neurotoxicity of AD AEVs and NEVs

As an initial test of the hypothesis that AD AEVs cause neurotoxicity and membrane disruption due to their surface cargo of complement components, we stripped AEVs of their surface protein epitopes by trypsination, which abolished their effect on neuronal viability (SI Appendix, Methods section; Fig. S4). Next, to assess whether MAC assembly *in situ* on the plasma membrane underlies the neurotoxicity of AD AEVs and NEVs we co-treated rat cortical neurons with CD59 and AEVs or NEVs from AD and control participants and found that CD59 abrogated the AD AEV- and NEV-mediated cell viability decrease on the MTT assay (Fig. 5a). [For NEVs, F for Condition (2, 74.636) = 3.687, p = 0.03; AD vs. Normal, p = 0.013; AD plus CD59 vs. Normal, p = 0.859; AD vs. AD plus CD59, p = 0.113. For AEVs, F for Condition (2, 90.658) = 7.818, p = 0.001; AD vs. Normal, p = 0.001; AD plus CD59 vs. Normal, p = 0.609; AD vs. AD plus CD59, p = 0.016.] This was further validated in a model of human induced Pluripotent Stem Cell-derived neurons subjected to the EthD-1 assay, the most stringent and relevant assay to the mechanism of complement and MAC-mediated cytotoxicity, in which co-treatment with CD59 blocked the AD AEV-mediated membrane disruption (AD vs. AD plus CD59 AEVs, p = 0.009; AD plus CD59 vs. Normal AEVs, p = 0.863) (Fig. 5b, c). Similar, but weaker, protective effects were produced by other endogenous inhibitors that bind and destabilize the C3 and C5 convertases upstream from MAC assembly: complement receptor 1 (CR1; also known as CD35), which inhibits the classical cascade (AD vs. AD plus CR1, p = 0.012; AD plus CR1 vs. Normal AEVs, p = 0.923), and Factor H (AD vs. AD plus Factor H, p = 0.011; AD plus Factor H vs. Normal AEVs, p = 0.698) and at trend-level the decay-accelerating factor (DAF; also known as CD55) (AD vs. AD plus DAF, p = 0.091; AD plus DAF vs. Normal AEVs, p = 0.414), which inhibit the alternative cascade. Importantly, effects of co-treatment with AD AEVs and only Factor I, which can degrade specific complement effectors only when aided by co-factors (e.g. Factor H and CR1) (7), were not different from effects of AD AEVs alone (AD vs. AD plus Factor I, p = 0.349; AD plus Factor I vs. Normal AEVs, p = 0.053).

**Figure 5.**
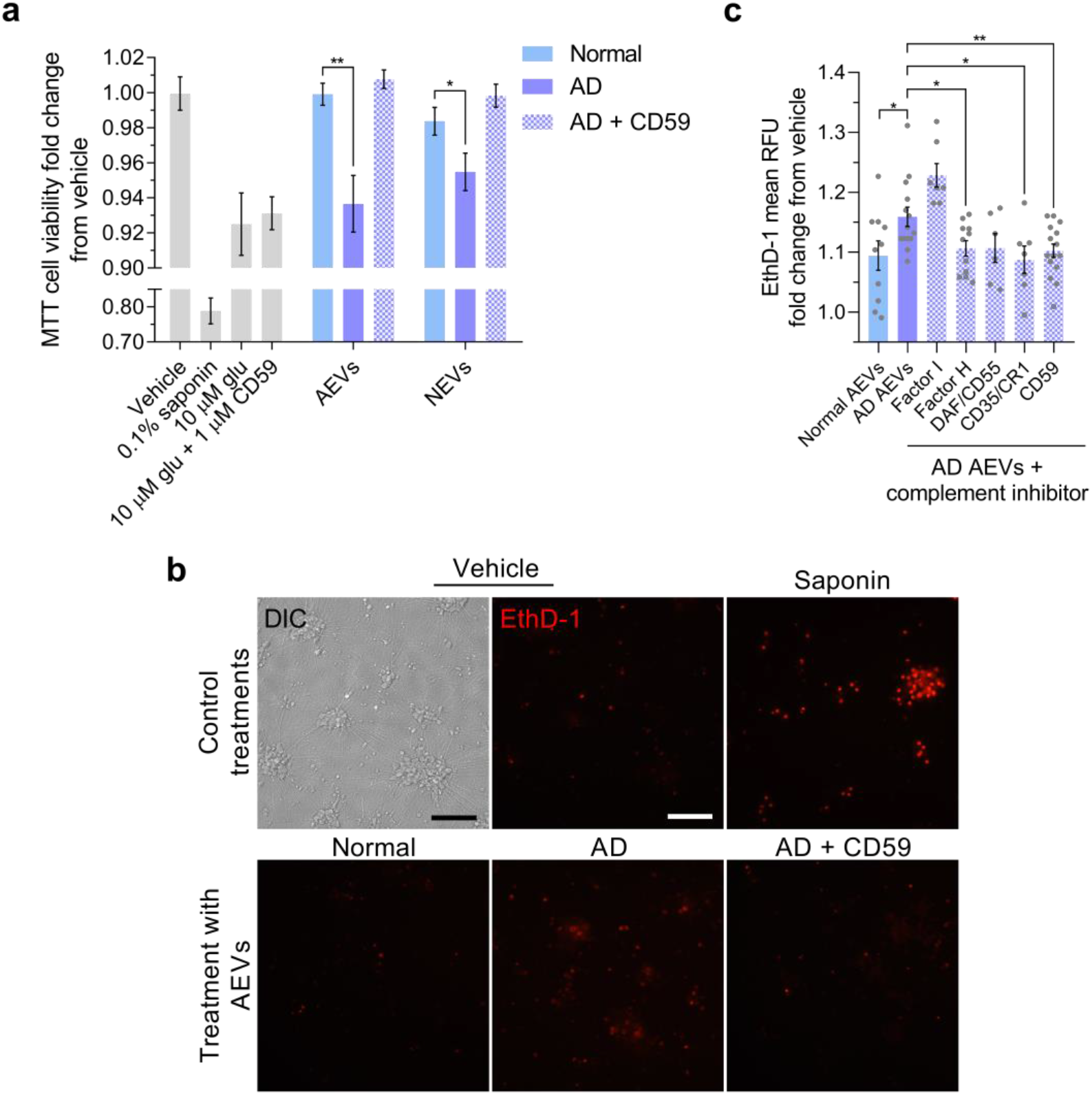
Complement inhibition prevents the neurotoxicity of AD AEVs and NEVs. **a,** Bar graph showing significant decreases in MTT cell viability (fold change from vehicle) observed in E18 rat cortical neurons treated with AEVs and NEVs from 4 AD participants compared to neurons treated with AEVs and NEVs from the same 4 AD participants co-incubated with MAC inhibitor CD59 and compared to neurons treated with AEVs and NEVs from 4 normal controls; 600 EVs/neuron for 48 hours were used. Neurons treated with 0.1% saponin detergent and 10 μM glutamate were used as positive controls for neurotoxicity. Co-treatment of neurons with 10 μM glutamate and 1 μM CD59 shows that CD59 does not exacerbate the glutamate-mediated cell viability impairment nor rescues it. Each bar represents the mean value ± SEM obtained from neurons treated in triplicate wells in two individual experiments. For NEVs, F for Condition (2, 74.636) = 3.687, p = 0.03; AD vs. Normal, p = 0.013; AD plus CD59 vs. Normal, p = 0.859; AD vs. AD plus CD59, p = 0.113. For AEVs, F for Condition (2, 90.658) = 7.818, p = 0.001; AD vs. Normal, p = 0.001; AD plus CD59 vs. Normal, p = 0.609; AD vs. AD plus CD59, p = 0.016. **b,** hiPSC-derived neurons incubated with AD AEVs at 800 EVs/mL for 48 hours show an increased nuclear EthD-1 fluorescence in comparison with neurons treated with AEVs from normal controls and vehicle-treated cells. Co-incubation with CD59 blocks the AD AEV-mediated EthD-1 fluorescence increase. Neurons treated with 0.1% saponin detergent were used as a positive control of neurotoxicity. Scale bar, 200 μm. **c,** Bar graph with individual values in grey showing the EthD-1 mean relative fluorescence units (RFU) fold change over vehicle ± SEM from hiPSC-derived neurons treated with AEVs from 5 AD participants with or without the indicated complement inhibitors and 5 normal controls in duplicate or triplicate. *p <0.05, **p <0.01, *****p<0.0001.

## Discussion

We demonstrated that AEVs and NEVs circulating in the blood of AD patients can be readily internalized by neurons, induce MAC deposition on their surface, cause membrane disruption and neurodegeneration, and decrease cell viability. These neurodegenerative effects were not produced by multicellular-origin CD81+ and total plasma EVs from AD patients or AEVs and NEVs from FTLD or cognitively normal participants and were blocked by the MAC inhibitor CD59 and additional endogenous complement inhibitors, suggesting that neuronal MAC assembly represents a necessary effector for AEV and NEV-mediated neurotoxicity in AD. Whereas previous studies have established the trans-neuronal spread of Aβ and tau via EVs (4, 34, 35), our study contributes to this growing field by demonstrating that EVs can also mediate the propagation of neuroinflammatory mediators in AD. These findings suggest a novel mechanism for neurodegeneration in AD, which involves neuronal internalization of AEVs and NEVs containing high levels of toxic complement components and low levels of endogenous inhibitors and recycling of their cargo to the plasma membrane leading to *in situ* complement activation and MAC assembly. An additional, albeit less likely, possibility, given protective effects of pre-treatment with CD59 and other complement inhibitors, is that preformed MAC, which we have shown to be present in AEVs (28), may be transferred to the plasma membrane of recipient neurons during EV fusion.

These conclusions are supported by employing multiple conditions and controls. To rule out an indiscriminate neurotoxic effect of circulating AEVs and NEVs, AD patients were compared to cognitively normal controls. To show neurodegenerative disease specificity for AD, we assessed FTLD AEV effects. To show specificity for neuronal and astrocytic EV origin and to rule out the involvement of some putative soluble factor contaminating our EV preparations, we tested multiple circulating EV types from the same subjects isolated through similar immunocapture procedures, specifically, L1CAM+ (neuronal), GLAST+ (astrocytic), CD81+ (variable cell types), as well as total EVs, showing that only neuronal and astrocytic EVs are neurotoxic. To assess the magnitude of EV-mediated toxicity we showed it to be comparable to neurotoxicity by 10 μM glutamate and sub-nanomolar oligomeric Aβ_42_, therefore physiologically relevant (31, 36). By showing that toxicity is abolished by EV trypsinization, we confirmed that it is mediated by surface proteins, such as molecular anchors that mediate EV uptake by recipient neurons (37) and/or complement components. To rule out that the human-origin of AD EVs may be responsible for toxicity to rat cortical neurons and confirm that identified mechanisms apply to human neurons, we also assessed hiPSC-derived neurons. Whereas the confluency achieved by hiPSC-derived neurons did not allow for assessment of toxicity with some of the assays conducted on rat cortical neurons (such as the MTT neuronal viability assay), we replicated the key findings of induced membrane disruption and its rescue by complement inhibitors using the EthD-1 assay. Moreover, the human subjects, from whom EVs were derived, were carefully selected (patients had high probability for AD based on CSF biomarkers (38) or met clinical criteria for two types of FTLD; controls have been followed longitudinally over many years and have not shown cognitive decline). Studying human circulating EVs rather than culture-derived EVs (such as in (39, 40)) renders greater validity to the findings. EVs from individual subjects were not pooled to avoid the possibility that neurotoxicity could be attributable to a single subject and respect their biological variability. A limitation of the study is the relatively small number of human subjects that were used, which was dictated by the need for homogeneity of the biological material used across multiple experiments, conditions and controls. Since our primary focus was to investigate mechanisms, biological variability was recognized and respected in the analysis, but not exhaustively studied. Future studies should investigate the functional and neurotoxic properties of AD patient EVs in larger cohorts and assess their potential clinical significance.

These findings are consistent with *in vitro* and *in vivo* studies showing that de-regulation of the brain complement system underlies synaptic loss in AD, which can be rescued by complement inhibition (6, 14), but critically contribute the role of EVs in the process. For years, the brain had been considered an immune-privileged organ. Today, it is known that various brain cells synthesize immune protein components of both the classical and alternative complement pathways including factors required for MAC assembly, essentially, a membrane pore-forming complex mediating cell death through osmotic lysis. Moreover, the MAC endogenous inhibitor CD59 is expressed by neurons (41) and astrocytes (41, 42), likely to protect them from uncontrolled neuroinflammation (43, 44). CD59 deficiency causes catastrophic neuroinflammatory conditions (45–47). In AD, protein levels of CD59 and other complement inhibitors are diminished, whereas those of complement effectors increased (44, 48), rendering neurons susceptible to complement attack (49). Here we show the effects of this attack, as increased membrane permeability (presumably the result of pore formation), necroptosis activation, neurodegeneration with loss of neurites and decreased cell viability.

Recent evidence challenges the notion that neuronal loss is universally detrimental in AD, since removal of damaged neurons may improve neuronal circuit function and functional outcomes (50). This raises the possibility that complement-mediated neuroinflammation may initially be a compensatory mechanism in the face of AD pathology by removing misfolded proteins, cellular debris, and inactive synapses, and enhancing plasticity, neural repair, and neuroprotection (51). However, *in vitro* and animal model studies show that persistent inflammation in AD becomes maladaptive resulting in neurotoxicity and synapse loss (2, 6, 52–54).

Neuronal death in neurodegenerative diseases may occur via apoptosis (a controlled process orchestrated by caspases), necrosis (uncontrolled cell death involving membrane disruption, release of cellular contents and inflammatory responses, not typically involving caspase activation) and, as recently recognized, necroptosis (a mode of programmed cell death involving membrane disruption, but not caspase activation) (55, 56). In accordance with *in vitro, in situ* and post-mortem AD studies showing that necroptosis is implicated in neuronal death in AD (57), we demonstrated that AEV-mediated neurotoxicity proceeds through membrane disruption in association with the phosphorylation of MLKL.

These findings motivate a set of intriguing hypotheses about AD pathogenesis and open new research avenues. EVs can be transported bidirectionally across the BBB (58) and inflammation can upregulate this process (59). The neuroinflammation of AD may enhance this bidirectional EV transport, which may result in AEVs and NEVs seeding AD pathology and inflammatory mediators to unaffected parts of the brain not only across neuroanatomical connections, but also via the systemic circulation. Additionally, circulating AEVs and NEVs of AD patients may be primed to uptake circulating complement factors and bring them back to the brain aggravating neuroinflammation. Future *in vivo* studies are required to elucidate whether passing through the systemic circulation is required for MAC- and EV-mediated neurotoxicity or whether brain cells are sufficient to provide all required components for MAC-induced neurotoxicity. Such experiments may include injection of AD patients’ EVs in the systemic circulation of animals to test for neurotoxicity and its reversal by systemic as opposed to intracerebral administration of complement inhibitors.

Although the amyloid hypothesis has been the predominant theory for AD pathogenesis (60), efforts aimed at translating it into effective therapies have failed, raising renewed interest in alternative therapeutic targets, particularly neuroinflammation. Our results suggest that inhibition of neurotoxic complement components and complexes or of their trans-cellular transfer through EVs may be therapeutically beneficial in AD. Complement inhibitors are already being developed for various clinical applications: for example, eculizumab is a well-tolerated monoclonal antibody that inhibits C5 and is being used in paroxysmal nocturnal hemoglobinuria (61), whereas a monoclonal antibody against C1q is well-tolerated in healthy volunteers and is being subjected to a phase I clinical trial (Clinicaltrials.gov number: NCT03010046). C3 inhibitors have been tested in non-human primate models of periodontitis and hemodialysis inflammation, with promising results (62, 63). Furthermore, oral administration of a C5 receptor inhibitor decreased pathology in AD mice (15). However, there are fundamental drawbacks for using systematically administered inhibitors in AD, since they may not easily cross the BBB and may also result in immunosuppression. Endogenous and/or engineered EVs may be used to overcome the BBB (64) and effectively deliver naturally occurring and/or synthetic complement inhibitors. The present study demonstrates that circulating AEVs and NEVs are readily uptaken and produce robust effects, motivating the use of modified AEVs and NEVs in brain-targeting therapies for AD.

## Materials and methods

### Human subjects

All participants donated blood as part of their participation in clinical studies at the National Institute on Aging (NIA) approved by the National Institutes of Health Institutional Review Board. All participants provided written informed consent. Procedures for sample collection and processing were identical for all samples. We analyzed samples from 7 participants with early AD and 6 age- and sex-matched cognitively normal controls randomly selected from the cohort used for the original report of high AEV complement in AD (28), as well as 2 participants with FTLD (one with behavioral variant Frontotemporal Dementia and one with Semantic Dementia) (Table 1-1). EVs isolated from individual subjects were not pooled and their effects were assessed and analyzed separately to respect their biological variability. AD participants had amnestic MCI or mild dementia (Clinical Dementia Rating (CDR) global score 0.5 or 1 respectively) with high probability AD by NIA–Alzheimer’s Association and International Working Group-2 criteria based on abnormal cerebrospinal fluid (CSF) levels of Aβ_1-42_, total tau and p181-tau (38, 65, 66); FTLD participants met clinical criteria (67). Normal controls were participants at the Baltimore Longitudinal Study on Aging conducted at the NIA who have remained cognitively normal for the course of their participation; they have Blessed Information Memory Concentration Test score < 4, and CDR = 0.

### Cell cultures

Primary cortical neurons were derived from Sprague Dawley^®^ rats at embryonic day 18 isolated from timed pregnant animals (Charles River Laboratories, Frederick, MD) and maintained as described previously (68). All procedures were approved by the National Institute on Aging Animal Care and Use Committee and complied with NIH guidelines. Briefly, the concentration of dissociated neurons in B27-supplemented neurobasal medium (Thermo Fisher Scientific) was determined using an hemocytometer and cells were cultured in polyethylenimine-coated plates and coverglass in a cell culture incubator at 37 °C and 5% CO_2_ for 14-21 days *in vitro* (DIV) prior to experiments.

### Ngn2-mediated neuronal differentiation of human iPSCs

Human iPSCs were maintained as feeder-free cells in Essential 8 medium on vitronectin-coated plates (all from Gibco, Carlsbad, CA) (69). The medium was changed every day and cells were passaged at 80%–90% confluency using TrypLE Express Enzyme (Gibco) supplemented with 10 μM Y-27632 (Stem Cell Technologies, Vancouver, Canada). The human iPSC line used in this study was the BC1 reprogrammed from peripheral blood mononuclear cells of a healthy male collected under a Johns Hopkins Institutional Review Board-approved clinical study (69).

The protocol followed for the neuronal differentiation of human iPSCs was adopted from Zhang et al. (70) who reported that forced expression of the single transcription factor Ngn2 can convert iPSCs into functional neurons with very high yield in less than 3 weeks. Briefly, human BC1 iPSCs were treated with Accutase (Innovative Cell Technologies, San Diego, CA) and plated as dissociated cells (25 × 10^4^ cells/well) on day −2 in a 6-well Matrigel-coated plate (BD Biosciences, San Jose, CA). On day −1, Ngn2 lentiviral infection (CHOP Research Vector Core, Philadelphia, PA) was performed using Polybrene (1 μg/μL; Sigma Aldrich, St. Louis, MO). The virus-infected cells were expanded, and frozen stocks made for future differentiation experiments. TetO/NGN2 gene expression was induced by Doxycycline (2 μg/mL; Clontech, Madison, WI) on day 0. Puromycin (2.5 μg/mL) was added to the medium on day 1 for 24 hours. Surviving cells were harvested on day 2 and plated on a Matrigel-coated 24 well plate at a concentration of 1 x 10^5^ cells/well. Cells were fed with neural differentiation media containing B27, BDNF, NT3 every other day until day 12. Cells were treated with 2 μM cytosine β-D-arabinofuranoside hydrochloride (Ara-C; Sigma Aldrich) on day 4 to reduce the proliferation of non-neuronal cells. Doxycycline was discontinued after day 12 and cells were fed every two days thereafter until day 21 when the neurons were mature enough to harvest.

### Isolation of astrocyte- and neuronal-derived extracellular vesicles from human plasma

All blood draws were conducted between 7 and 10 am and after an overnight fast at the NIA Clinical Unit following standard procedures. Approximately 10 mL of venous blood were collected in plasma separator tubes containing EDTA, incubated for 10 minutes at room temperature (RT) and then centrifuged at 3000 rpm for 15 minutes at RT. Supernatant plasma was divided into 0.5 mL aliquots and stored at −80 °C until further use. Hemolysis was ruled out using spectrophotometry (data not shown). Pre-analytical factors for blood collection and storage comply with guidelines for EV biomarkers (71, 72).

Plasma samples were thawed on ice and immediately subjected to isolation of AEVs and NEVs using a methodology extensively described elsewhere (25, 26, 28). Briefly, fibrinogen and fibrin proteins known to impede efficient EV recovery were removed using 5 U/mL of thrombin (System Biosciences, Inc.; Mountainview, CA) for 30 minutes at RT followed by addition of 495 μL of Dulbecco’s PBS-1X (DPBS) supplemented with protease and phosphatase inhibitors and centrifugation at 6,000 *g* for 20 minutes at 4 °C. The supernatant was transferred to a sterile 1.5 mL microtube and total EVs were sedimented by incubation with 252 μL of ExoQuick^®^ (System Biosciences) for 1 hour at 4 °C followed by centrifugation at 1,500 *g* for 20 minutes at 4 °C. The EV-depleted supernatant was transferred to a sterile 1.5 mL microtube and stored in −80 °C. Pelleted total EVs were resuspended by overnight gentle rotation mixing at 4 °C in 500 μL of DPBS supplemented with protease and phosphatase inhibitors. Resuspended total EVs were incubated with 4 μg of anti-GLAST (i.e. antibody against the astrocyte cell surface antigen-1; RRID: AB_2733473; Miltenyi Biotec, Auburn, CA), anti-human CD171 (i.e. antibody against neural cell adhesion molecule L1CAM; RRID: AB_2043813; Thermo Fisher Scientific, Waltham, MA) or anti-human tetraspanin CD81 (Ancell Corporation, Bayport, MN) biotinylated antibodies, to immunocapture AEVs, NEVs and endosome-derived CD81+ EVs of variable cell origin (73) respectively, for 4 hours at RT. EV-antibody complexes were incubated with 30 μL of washed Pierce™ Streptavidin Plus UltraLink™ Resin (Thermo Fisher Scientific) for 1 hour at RT; EV-antibody-bead complexes were allowed to sediment by gravity followed by removal of unbound EVs and soluble proteins in the supernatant. Bound AEVs, NEVs and CD81+ EVs were eluted using 200 μL of 0.1 M glycine followed by the immediate transfer of the supernatant to a sterile microtube containing 20 μL of 1 M tris buffer for pH neutralization. 10 μL of intact EVs were used for determination of particle concentration and diameter using nanoparticle tracking analysis (NTA) (Nanosight NS500; Malvern, Amesbury, UK). The remaining 210 μL of intact EVs was aliquoted into sterile microtubes in 20 μL aliquots or mixed with 1.5 parts of MPER lysis buffer (Thermo Fisher Scientific) supplemented with protease and phosphatase inhibitors for downstream immunoassays, and then stored at −80 °C until further use.

The size and morphology of intact AEVs were assessed using Transmission Electron Microscopy. EVs were absorbed on to a 400-mesh carbon coated grid (Electron Microscopy Sciences, Hatfield, PA) for 1 minute and then quickly rinsed using ddH2O followed by staining with uranyl acetate. All imaging was done on a Zeiss Libra 120 (Zeiss, Thornwood, NY) with an Olympus Veleta camera (Olympus America, Center Valley, PA).

### Characterization of AEVs and NEVs by immunoblotting and ELISA

Sequential EV purification through the aforementioned two-step EV immunoprecipitation protocol was confirmed by assessing positive and negative EV protein markers in EV-depleted plasma, total EV isolates and immunoprecipitated AEVs, NEVs and CD81 + EVs using immunoblotting. The protein concentration was determined using the Bradford protein assay (Biorad, Hercules, CA). One μg of total protein per sample was resolved by sodium dodecyl sulfate polyacrylamide gel electrophoresis (SDS-PAGE) using 4-12% bis-tris gels in MOPS SDS running buffer (NuPAGE^®^ Novex^®^ SDS-PAGE system; Thermo Fisher Scientific) and transferred to polyvinylidene fluoride membranes (iBlot^®^ 2 gel transfer system; Thermo Fisher Scientific). Membranes were blocked using the Odyssey^®^ blocking buffer in TBS (Licor Biosciences, Lincoln, NE) for 1 hour at RT and incubated overnight at 4 °C with fluorescently labelled antibodies (IRDye^®^ 680RD antibody infrared dye; Licor Biosciences) targeting apolipoprotein A1 (0.4 μg/mL; RRID: AB_2242717, cat. no. AF3664, R&D Systems, Minneapolis, MN) as an indicator of lipoproteins, the cis-golgi marker GM-130 (0.2 μg/mL; RRID: AB_880266, cat. no. 52649, Abcam, Cambridge, MA) used as a negative EV marker, and the membrane and intra-vesicular positive EV markers CD81 (1:500; cat. no. EXOAB-CD81A-1, System Biosciences) and alix (0.4 μg/mL; RRID: AB_11023702, cat. no. NBP1-90201, Novus Biologicals, Littleton, CO), respectively (29). Antibody excess was washed five times with tris buffer saline supplemented with 0.05% tween-20 detergent for 5 minutes and blots were scanned using the Odyssey^®^ CLx imaging system (Licor Biosciences).

### Neurotoxicity assays

The effects of AEVs, NEVs, CD81+ and total EVs, from normal control, AD and FTLD participants, on the metabolic activity of primary neurons were evaluated using the tetrazolium salt 3-(4,5-dimethylthiazol-2-yl)-2,5-diphenyltetrazolium bromide (MTT) assay (Trevigen, Gaithersburg, MD), a colorimetric test based on the enzymatic activity of NADPH-dependent cytoplasmic oxidoreductase enzymes that catalyze the reduction of the membrane permeable MTT into colorimetric formazan products. Additionally, EV effects on neuronal membrane integrity were assessed using the ethidium homodimer (EthD-1) assay (Cell Biolabs, San Diego, CA), which is based on the detection of the membrane impermeable EthD-1 dye which fluoresces when bound to DNA. The EV concentration used to treat neurons and treatment duration were based on a previous study, in which the neurotoxicity of CSF EVs of AD patients was evaluated using E18 rat cortical neurons (30), and confirmed by experiments assessing the optimum concentration- and time-dependency of neurotoxicity of immunoprecipitated EVs.

E18 rat cortical neurons cultured for 14-21 DIV in PEI-coated 96-well plates at a confluency of 100,000 neurons per well in 200 μL of neurobasal media were treated with AEVs, NEVs, CD81+ and total EVs of AD, FTD and Normal participants at 50, 100, and 600 EVs/neuron in 20 μL of neurobasal media for 24, 48 and 72 hours in triplicate and then subjected to MTT and EthD-1 assays following the manufacturers’ instructions. Neuronal culture in corner wells of the plate was avoided and 300 μL of ultrapure distilled water added instead to prevent artificial alterations in cell viability caused by medium evaporation, a phenomenon known as the ‘edge effect’. Treatments with 10 and 100 μM glutamate for 24 hours and 0.1% saponin detergent for 10 minutes were used as positive controls of neurotoxicity whereas neurons treated with 20 μL of neurobasal media (vehicle treatment) were used as a negative control. Plates were read using the multifunctional microplate reader Synergy^™^ H1 and data collected using the Gen5^™^ microplate data collection and analysis software (BioTek Instruments). Given that the fluorescent signal from EthD-1 plates is emitted from adherent cells unevenly distributed in the bottom of wells, the surface area of each well was scanned using the 9×9 area scan read module and the average signal used for data analysis. MTT and EthD-1 signals from EV-treated samples and positive controls of neurotoxicity were normalized to vehicle treatment prior to statistical analysis. To evaluate if the neurotoxicity of AEVs and NEVs on the MTT assay depends upon MAC formation, the MTT assay was carried with co-treatment with CD59. Primary neurons and EVs were pre-incubated with active recombinant human CD59 (cat. no. 1987-CD, R&D Systems) at a final concentration of 1 μM (stock at 10 μM) for 2 hours in the cell culture incubator prior to treatment of cells with EVs.

The complement-mediated neurotoxicity of AD AEVs was further validated in a human-based model using hiPSC-derived neurons cultured in Matrigel-coated 12-well plates for 12-18 DIV at a cell confluency of 100,000 neurons per well in a final volume of 500 μL of neural differentiation media. The EthD-1 membrane disruption assay was carried out using hiPSC-derived neurons treated with 800 EVs/neuron in 35 μL of neurobasal medium (final concentration of EVs in culture: 1.5 x 10^8^ EVs/mL) for 48 hours with and without the addition of inhibitors of the classical pathway (Factor I and CR1), alternative pathway (Factor H and DAF) and terminal pathway (CD59). The concentration used for each inhibitor was determined using the reported EC50 values, as follows: Factor I, 70 nM, stock at 2 μM; CR1, 10 nM, stock at 1.88 μM; Factor H, 235 nM, stock at 10 μM; DAF, 70 nM, stock at 2 μM; CD59, 500 nM, stock at 10 μM. CR1 (cat. no. 5748-CD), Factor H (cat. no. 4779-FH), DAF (cat. no. 2009-CD) and CD59 (cat. no. 1987-CD) were acquired from R&D Systems, whereas Factor I (cat. no. 00003426-Q01) was acquired from Novus Biologicals. EthD-1 results acquired using the fluorescence plate reader were further validated qualitatively by visualization of EthD-1 positive nuclei using an Olympus IX50 inverted microscope equipped with a U-LH100HG fluorescence light source (Olympus America) and an AxioCam HrC camera (Zeiss).

### Assessment of neurodegeneration by immunocytochemistry

The neurite density of EV-treated rat cortical neurons was assessed by means of β III-tubulin (neuron-specific class III tubulin, also known as Tuj-1) fluorescent immunoreactivity to evaluate whether EV treatments induce neurodegeneration. Neurons cultured in PEI-coated 8-well chambered borosilicate coverglass (Thermo Fisher Scientific) at a confluency of 200,000 neurons/well in 500 μL of neurobasal media were treated in duplicate with 20 μL of AEVs, NEVs and CD81+ EVs from AD and normal participants in neurobasal media at 600 EVs/neuron (final concentration of EVs in culture: 2.4 x 10^8^ EVs/mL) for 48 hours using 100 μM glutamate treatment as a positive control of neurotoxicity. After treatment with EVs, neurons were fixed with 4% FA followed by membrane permeabilization with 0.1% triton X-100 detergent in DPBS for 15 minutes at room temperature. Excess detergent was washed three times with DPBS and non-specific antibody binding was blocked with DPBS supplemented with 2% bovine serum albumin for 1 hour at RT. Then, cells were incubated with 1 μg/mL of mouse anti-β III-tubulin antibody (RRID: AB_2256751, cat. no. 78078, Abcam) in blocking solution overnight at 4 °C, followed by five washes with DPBS and incubation with 1 μg/mL Alexa Fluor^™^ 647 donkey anti-mouse antibody (RRID: AB_162542, cat. no. A-31571, Thermo Fisher Scientific) for 1 hour at RT. Excess of secondary fluorescent antibody was washed five times with DPBS and mounting media supplemented with the nucleic acid stain DAPI added prior to visualization of cells using fluorescence confocal microscopy. Ten to fifteen images per sample were acquired at 10X and 25X magnifications and analyzed using the image processing software Fiji (74). The ‘moments’ threshold image filter was applied and the surface image area of β III-tubulin selected, measured and normalized to the amount of DAPI+ nuclei counted using the ImageJ Image-based Tool for Counting Nuclei (ITCN) plugin (Center for Bio-Image Informatics, University of California, Santa Barbara).

MAC immunocytochemistry was conducted in EV-treated rat cortical neurons to evaluate MAC expression in recipient cells. Neurons were treated with AEVs from AD and normal participants for 1 hour and then subjected to fluorescent immunolabelling for MAC using a mouse anti-human C5b-9 antibody (1 μg/mL; RRID: AB_2067162, cat. no. M0777, Dako, Carpinteria, CA) targeting an epitope in the poly-C9 component of MAC in association with C5b, a protein quaternary structure characterizing the active complex (75), and showing cross-reactivity with the C5b-9 equivalent protein in rat (76). To determine the neuronal localization of MAC immunoreactivity, cells were co-labelled with rabbit anti-β III-tubulin antibody (RRID: AB_444319, cat. no. 18207, Abcam) at 1 μg/mL using Alexa Fluor^™^ 488 goat anti-mouse and 647 goat anti-rabbit (RRID: AB_2534069, cat. no. A11001; and RRID: AB_2535813, cat. no. A21245; Thermo Fisher Scientific) as secondary antibodies. MAC and β III-tubulin immunoreactivity were visualized via fluorescence confocal microscopy and 5-10 images per sample acquired at 10, 20 and 63X magnifications. Images were analyzed using the image processing software Fiji and the mean fluorescence of MAC positive regions of interest selected using the ‘moments’ threshold image filter determined.

### Assessment of EV-mediated activation of apoptosis and necroptosis

To determine if apoptosis and/or necroptosis underlie the neurotoxicity of AD AEVs, E18 rat cortical neurons cultured at a density of 1.6 × 10 ^6^ neurons/well in PEI-coated 12-well plates were treated with 100 μL of AEVs from AD and normal participants in neurobasal media at a final concentration of 600 EVs/neuron (9.6 × 10 ^8^ EVs/mL) for 48 hours in triplicate. After incubation, culture media was removed and neurons were washed once with DPBS followed by cell lysis using 50 μL of RIPA buffer (Thermo Fisher Scientific) supplemented with phosphatase and protease inhibitors. The protein concentration of pooled triplicates was determined using the BCA protein assay (Thermo Fisher Scientific) and 8 μg of total protein subjected to immunoblotting against cleaved caspase-3 and −7 (1:200 dilution; RRID: AB_2341188 and RRID: AB_11178377; cat. no. 9661S and 8438S, Cell Signaling, Danvers, MA) as well as phosphorylated (1:500 dilution; cat. no. MABC1158, Millipore Sigma, Burlington, MA) and total MLKL (1:500 dilution; RRID: AB_2737025, cat. no. 172868, Abcam) using β-actin (RRID: AB_306371, cat. no. 8226, Abcam) as loading control, using 10% bis-tris gels in MES SDS running buffer.

The EV-mediated activation of apoptosis was also assessed using an in-vivo fluorescence microscopy assay designed to detect the abundance of active caspases in live cells (Image-iT^™^ Live Green Poly Caspases Detection Kit, cat. no. I35104, Thermo Fisher Scientific). This assay is based on the cell permeant and noncytotoxic FLICA^™^ fluorescent inhibitor of caspases that binds covalently to the enzymatic reactive center of activated caspases, thus serving as a reporter of active caspases in EV-treated neurons. E18 rat cortical neurons cultured for 14-21 DIV in PEI-coated 8-well chambered cover glasses at a confluency of 100 neurons/well in 500 μL of neurobasal media were treated with 20 μL of AEVs and NEVs in neurobasal media at 600 EVs/neuron in duplicate for 48 hours using 0.1% saponin as a positive control for neurotoxicity. After treatment with EVs, neurons were subjected to the *in vivo* poly-caspases detection assay according to the manufacturer’s instructions and visualized under a fluorescence confocal microscope. 3-10 images per sample were acquired at 10X magnification and the mean fluorescence of FLICA positive regions of interest was determined using the Fiji’s ‘max entropy’ threshold image filter. Signals from EV- and saponin-treated samples were normalized to vehicle treatment and subjected to statistical analysis.

### Experimental design and statistical analyses

Analyses were performed using SPSS version 25. For analyses of biological effects of EV treatments, we used mixed linear models for repeated measures to assess the fixed effects of factors Group (i.e. AD, normal, and FTLD), on vehicle-normalized values of MTT, EthD-1 and neurite density assays. To appropriately handle biological and technical variability respectively, we analysed individual measurements, modelling correlated random effects for Subject and using Well (from 96-well plates of cultured neurons) and Plate as repeated measures (15 well repetitions on average per EV type per Subject). For significant effects of Group, we conducted t-tests of least square means to determine the direction and significance of pairwise differences (e.g. comparing effects of AEVs from AD vs. normal participants); reported comparisons were Bonferroni-corrected. Alternative models including participant age and sex did not result in better fit (by AIC and BIC criteria). Mixed models for repeated measures were also used to analyse MTT and EthD-1 rescue experiments assessing the effect of Condition (AEVs alone, AEVs plus CD59, AEVs plus Factor H, AEVs plus Factor I, AEVs plus DAF). Models comparing EV types (AEVs, NEVs, CD81+ and total EVs), included EV type as a Factor and the Group*EV type interaction term. Comparison of results from additional experiments (i.e. concentration- and time-dependent EV treatments, C5b-9 accumulation and activation of apoptosis and necroptosis in EV-recipient neurons, CD59 protein levels in EVs using ELISA and neurotoxicity of trypsinized vs. intact EVs) were made using two-tailed unpaired t-tests or ordinary one-way ANOVA corrected for multiple comparisons using the Dunnet test. Statistical details of experiments can be found in the figure legends and Results.

## Acknowledgements

This research was supported in part by the Intramural Research Program of the National Institute on Aging, National Institutes of Health and grants from the Maryland Stem Cell Research Fund. We would like to thank Dr. Erez Eitan and Mr. Sahil Chawla for their intellectual and practical assistance with experiments.

## Supplementary information

### Supplementary methods

#### Assessment of CD59 protein levels in EVs by ELISA

The abundance of the endogenous complement inhibitor CD59 in undiluted MPER-lysed AEVs, NEVs and CD81+ EVs of AD patients and normal controls was evaluated using ELISA (cat. no. ELH-CD59, RayBiotech, Norcross, GA). The protein levels of CD81 were used as a normalization factor for EV load (cat. no. CSB-EL004960HU, Cusabio Technology, College Park, MD). ELISA plates were read using the multifunctional microplate reader Synergy™ H1 and data collected using the Gen5^™^ microplate data collection and analysis software (BioTek Instruments, Winooski, VT). Absorbance signals above the limit of detection (LoD) were not extrapolated to protein concentration as they were below the lowest limit of quantification (LLoQ) and are reported as raw signal. The LoD was defined as the mean signal of the diluent blank plus 2.5 times its standard deviation whereas the LLoQ was determined by the standard solution with signal above the LoD, CV among duplicates lower than 20%, and recovery between 80% and 120%.

#### Assessment of EV uptake by neurons

To evaluate the interaction of AEVs with recipient neurons, E18 rat cortical neurons at 21 DIV were treated with fluorescently labelled AEVs which were then tracked using fluorescence confocal microscopy to elucidate their sub-cellular localization. AEVs isolated from AD and normal participants were labelled with the red fluorescent lipid analogue PKH26 (Sigma Aldrich, St. Louis, MO) as instructed by the manufacturer with modifications. 100 μL of AEVs at a concentration of 10^10^ EVs/mL (10^9^ particles) by NTA were incubated with 5 μL of a 4 μM PKH26 solution in buffer C for 1 min at RT (buffer C lacks physiological salts and thus reduces the formation of dye micelles that could result in a false positive neuronal staining) (1). To control for the possibility that salts in the EV vehicle, i.e. DPBS, could induce the formation of dye micelles and thus interfere with the interpretation of results, we subjected DPBS alone to PKH26 using the same staining procedure as AEVs. To further minimize the potential for micelle formation, after EV labelling, 400 μL of DPBS were added to the solution followed by the removal of free dye using size-exclusion chromatography (Zeba^™^ spin desalting column; cat. no. 89892, Thermo Fisher Scientific). The concentration of PKH26+ AEVs was assessed by NTA and 300 μL were added to E18 rat cortical neurons cultured for 21 DIV in PEI-coated wells of a borosilicate 8-well chambered coverglass (Thermo Fisher Scientific) at a concentration of 200,000 neurons per well in a final volume of 500 μL of neurobasal media (EV concentration in culture: 3-8 × 10^9^ EVs/mL). Neurons were incubated with PKH26+ AEVs for 1 hour, washed two times with DPBS and fixed with 4% formaldehyde (FA) in DPBS for 10 minutes at RT. Excess FA was washed three times with DPBS. Mounting media supplemented with the nucleic acid stain DAPI (cat. no. P36962, Thermo Fisher Scientific) was added prior to visualization of cells using an LSM 710 confocal laser scanning microscope (Zeiss).

#### Trypsination of extracellular vesicles

External EV protein epitopes were proteolytically cleaved using trypsin in order to evaluate whether neuronal uptake and EV-mediated neurotoxicity depend upon surface protein moieties. Trypsination of EVs was carried out using trypsin immobilized on beaded agarose (immobilized TPCK trypsin; Thermo Scientific, Inc.) to eliminate trypsin carryover to subsequent neuronal treatments and to avoid it causing EV permeabilization. Thirty μL of immunoprecipitated EV samples ranging from 1-10 × 10^10^ EVs/mL (0.5-1 mg/mL total protein) were incubated with 10 μL of washed trypsin bead slurry (recommended usage based on trypsin activity: 200 μL trypsin bead slurry per mg of total protein) for 30 minutes at 37 °C. Then, the trypsin gel was separated from the digestion mixture by centrifugation and trypsinized EVs in the supernatant were transferred to sterile microtubes. EV concentration and structure was assessed prior to neuronal treatments using NTA.

---

**Supplementary table 1.** Subjects demographics and clinical parameters. Variables except for sex are presented as mean ± standard deviation. P-values for differences were calculated using unpaired t-test. M = males; F = females. MMSE, Mini-Mental State Exam score; ADAS-cog, Alzheimer’s Disease Assessment Scale-cognitive subscale score; CDR, Clinical Dementia Rating scale.

|  | AD | Normal | FTLD | P-value (AD vs. Controls/<br>AD vs. FTLD) |
| --- | --- | --- | --- | --- |
| Number of subjects | 7 | 6 | 2 | --- |
| Age (years) | 73 ± 9 | 76 ± 7 | 78 ± 21 | .545/ .589 |
| Sex | 5F/2M | 4F/2M | 1F/1M |  |
| MMSE | 22 ± 5 | --- | 23 ± 9 | --- / .805 |
| ADAS-cog | 18 ± 5 |  | 20 ± 21 | --- / .744 |
| CDR global | .6 ± .2 | --- | .7 ± .3 | --- / .345 |
| CDR-sob | 3.2 ± 2 | --- | 3.5 ± 3 | --- / .880 |
| CSF Total Tau | 61 ± 12 | --- | 58 ± 33 | --- / .848 |
| CSF p181-Tau | 43 ± 13 | --- | 31 ± 13 | --- / .283 |
| CSF Aβ 1-42 | 168 ± 28 | --- | 237 ± 3 | --- / .014 |

**Supplementary figure 1.**
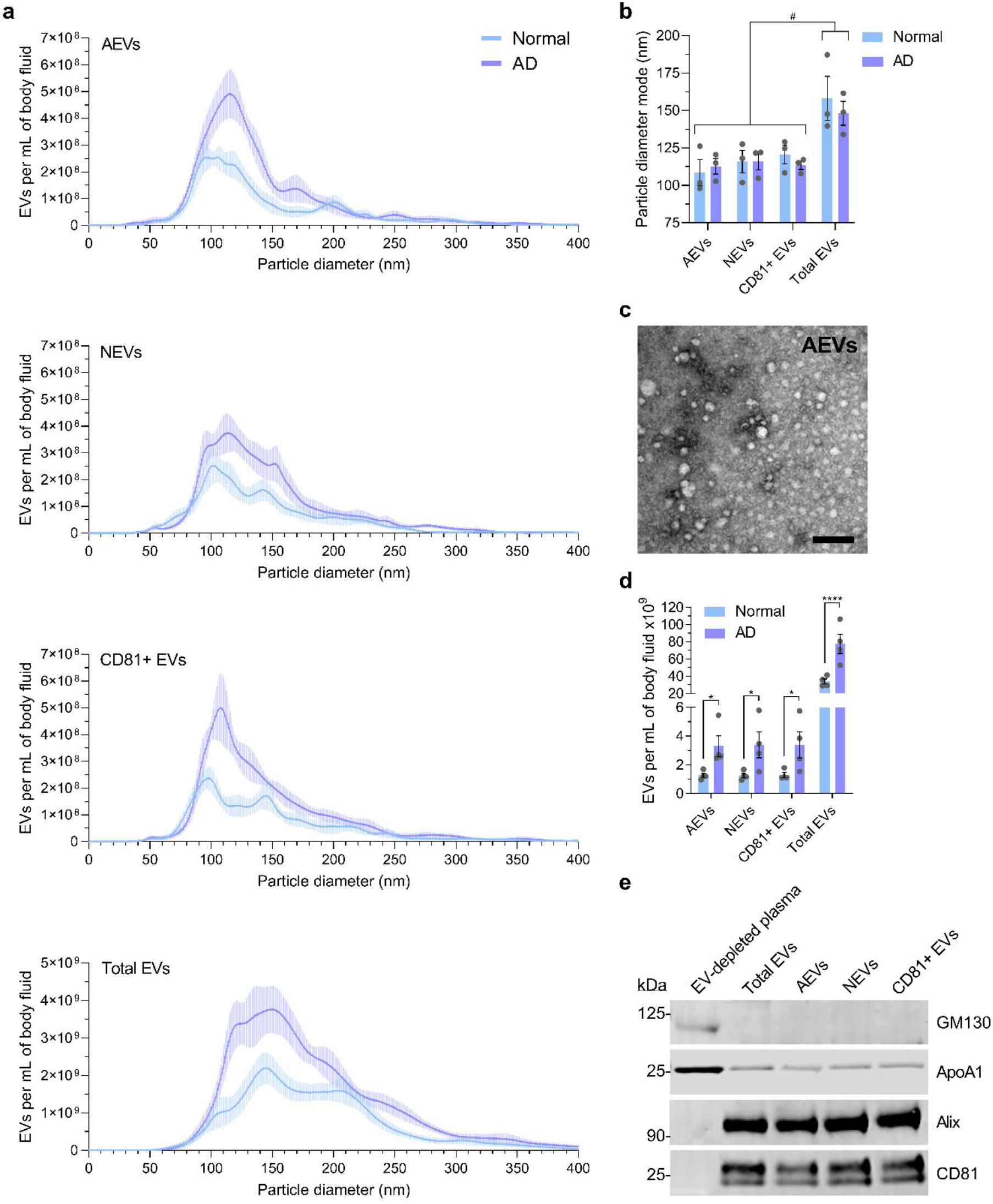
Characterization of extracellular vesicles. **a,** The graph presents EV concentration (particles/EVs per mL of plasma) as a function of particle diameter [determined using nanoparticle tracking analysis (NTA)] for immunoprecipitated AEVs, NEVs and CD81+ EVs, and total EVs isolated from the plasma of 3 AD patients and 3 normal controls. **b,** A bar graph showing the mean ± SEM of the particle diameter mode in **a** with individual values in grey. Statistical analysis: two-way ANOVA; significant differences indicated by the symbol ‘#’. **c,** Transmission Electron Microscopy negative staining of AEVs of a normal aging participant. Scale bar, 200 nm. **d,** A bar graph showing the mean ± SEM of the particle concentration in **a** with individual values in grey. Statistical analysis: two-way ANOVA of AD vs Normal; AEVs: *p = 0.0214, NEVs: *p = 0.0152, CD81+ EVs: *p = 0.0175, total EVs: ****p < 0.0001. Interestingly, all types of EVs from AD patients had higher particle concentrations compared to EVs from normal controls. **e,** Western blots showing the protein levels of Apolipoprotein A1 as an indicator of lipoproteins co-precipitated with EVs, the cis-Golgi marker GM-130 used as a negative EV marker, and the transmembrane and intra-vesicular positive EV markers, CD81 and alix respectively, in 1 μg of total protein from the EV-depleted plasma, total EVs, AEVs, NEVs and CD81 + EVs of a normal control participant.

**Supplementary figure 2.**
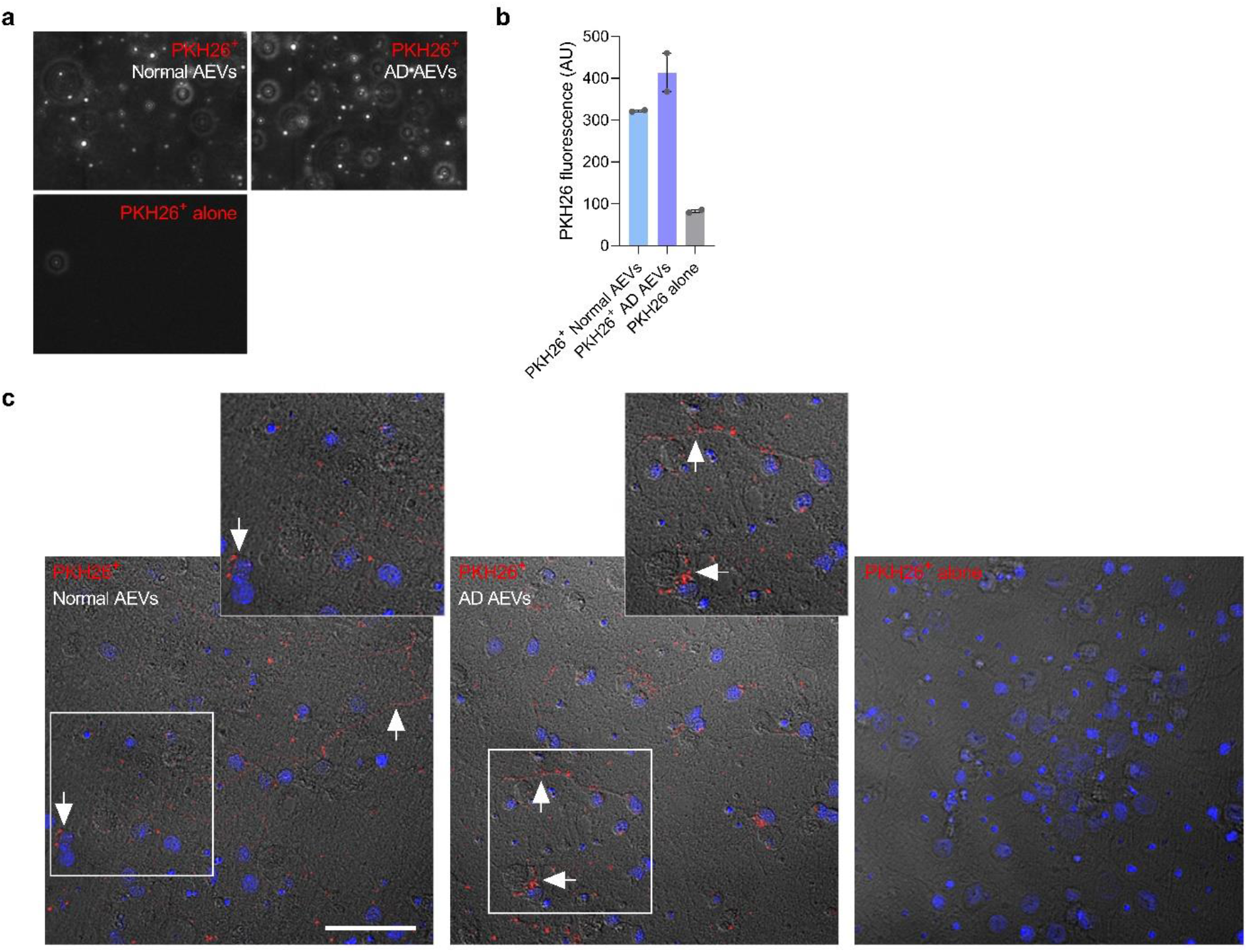
Neuronal internalization of AEVs. **a,** Nanoparticle tracking analysis video stills showing that the particles collected by size exclusion chromatography after labelling with PKH26 (PKH26+ AEVs from normal control and AD participants) are not PKH26 micelles (PKH26 alone). **b,** Mean PKH26 fluorescence intensity ± SEM of labelled AEVs from 2 AD patients and 2 normal controls compared to PKH26 alone confirm EV labelling. **c,** PKH26 (red) fluorescence microscopy images merged with its differential interface contrast (DIC) and DAPI nuclear counterstain (blue) of rat cortical neurons treated with PKH26+ AEVs from normal control and AD participants for 1 hour show the accumulation of AD and normal AEVs in the soma and neurites (arrows in image and magnified inserts). Treatment with PKH26 alone was used as a negative control. Scale bar, 200 μm.

**Supplementary figure 3.**
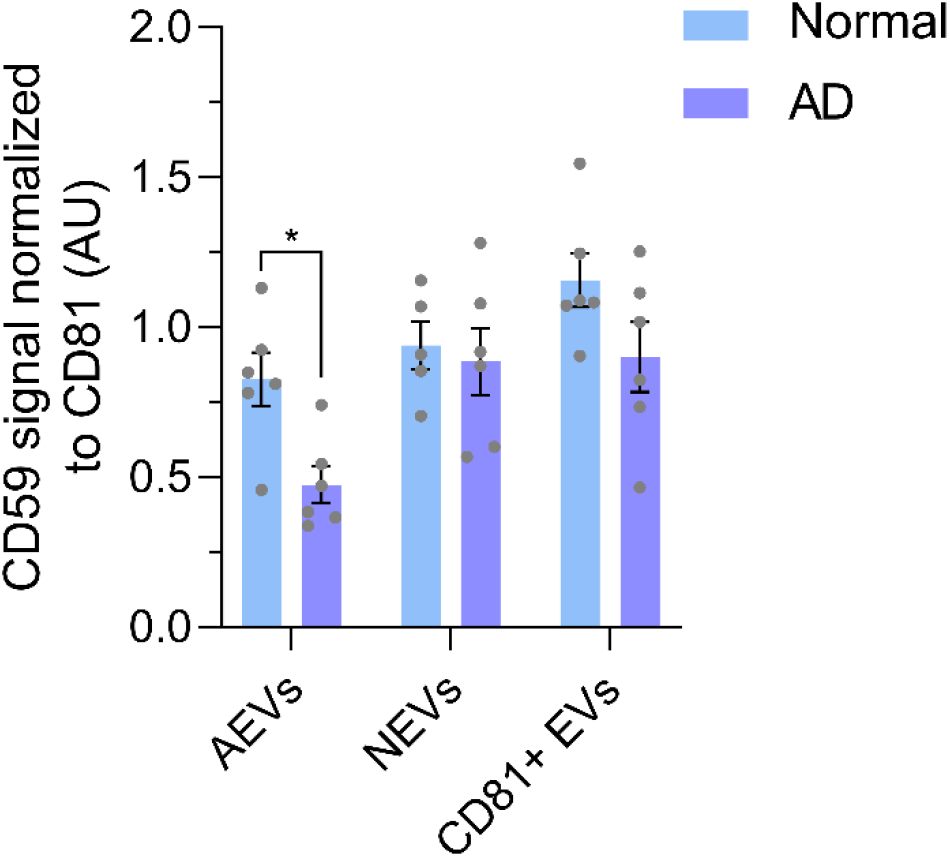
Decreased protein levels of the endogenous MAC inhibitor CD59 in AD AEVs. The protein levels of the endogenous MAC inhibitor CD59 was assessed in lysates of plasma AEVs, NEVs and CD81+ EVs of AD and normal control participants using ELISA. Each bar represents the mean value ± SEM of CD59 absorbance normalized to the EV input measured by CD81 ELISA from EV lysates of 6 AD subjects and 6 controls. Statistical analyses: two-way ANOVA of AD vs Normal; *p = 0.0360.

**Supplementary figure 4.**
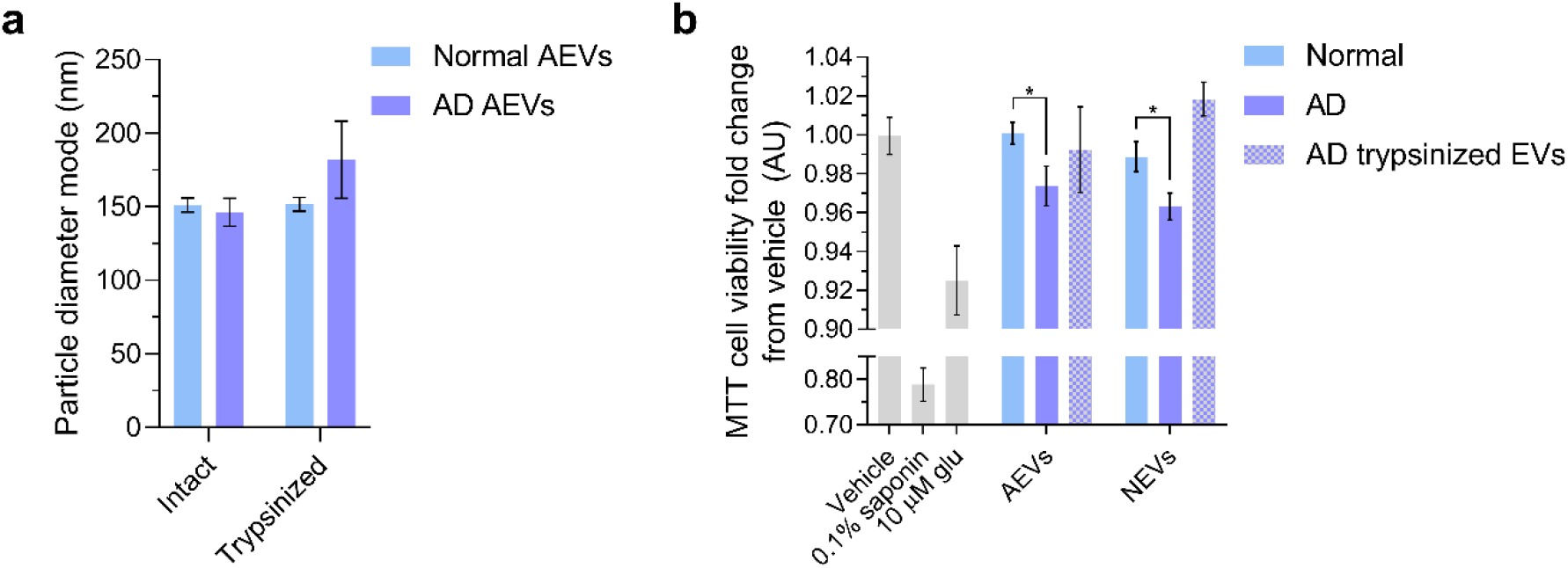
Trypsination of extracellular vesicles abolishes the neurotoxicity of AD AEVs and NEVs. **a,** A bar graph showing that the mean NTA particle diameter mode ± SEM of intact AEVs of 2 AD and 2 normal control participants does not change after trypsination, thus suggesting that digestion of surface tryptic peptides does not disrupt EVs. **b,** Bar graph showing that trypsination of AD AEVs and NEVs abolishes the significant decrease in MTT cell viability (fold change from vehicle) in E18 rat cortical neurons treated with AEVs and NEVs from 2 AD participants compared to neurons treated with AEVs and NEVs from 2 normal controls, respectively. Neurons treated with 0.1% saponin detergent and 10 μM glutamate were used as positive controls for neurotoxicity. Each bar represents the mean value ± SEM from triplicate wells and two different experiments. Analysis was based on multiple comparisons using one-way ANOVA: AD vs Normal AEVs, *p = 0.0238; AD vs Normal NEVs, *p = 0.0247.

